# High levels of antisense transcription from numerous genes in the obligate intracellular bacterium *Orientia tsutsugamushi*

**DOI:** 10.64898/2025.12.03.692040

**Authors:** Chitrasak Kullapanich, Regan Hayward, Lars Barquist, Jeanne Salje

## Abstract

*Orientia tsutsugamushi* (Ot) is a cytoplasmic, Gram-negative, obligate intracellular bacterium that causes the human disease scrub typhus. It has a highly repetitive genome whereby approximately 50% is composed of multiple copies of a proliferated integrative and conjugative element, the Rickettsial amplified Genetic Element (RAGE). Previous RNA sequencing analysis revealed a high level of antisense transcription in Ot, particularly from RAGE-encoded genes, with 18% of all genes having a sense:antisense ratio <1 and 5% with a ratio <0.1. In our current study, we have confirmed the earlier RNA sequencing findings via PCR-based methods and established that this antisense transcription is a consistent feature across eight distinct Ot strains. Furthermore, we have utilized PCR to monitor differences in sense and antisense expression between intracellular and extracellular bacteria. Our findings lend weight to the hypothesis that antisense transcription in Ot is a regulated mechanism, possibly playing a crucial role in the pathogen’s lifecycle during infection.

## Introduction

*Orientia tsutsugamushi* (Order Rickettsiales, Family Rickettsiaceae) (Ot) is an obligate intracellular Gram negative alphaproteobacterium that causes the mite-borne human disease scrub typhus^1,2^. This is endemic in many parts of rural Asia, with related species causing scrub typhus like illness recently reported from Latin America^3,4^ and the Middle East^5^. Symptoms begin 7-10 days after being bitten by an infected mite, and include headache, fever, rash, and a distinctive lesion called an eschar at the site of inoculation. If not treated with timely administration of effective antibiotics the infection can escalate to cause multiple organ failure and death, with a median mortality of 6% in untreated cases^6^.

Ot is an obligate intracellular bacterium that escapes from the endolysosomal pathway shortly after cellular internalisation and carries out its growth and replication directly in the eukaryotic cytoplasm^7,8^. It traffics to the perinuclear region using dynein-dependent motility^9^ mediated by the bacterial surface autotransporter protein ScaC^10^, where it undergoes replication within a tightly packed bacterial microcolony. Following replication, Ot exits infected cells using a budding mechanism, in which it buds off the surface encased in host plasma membrane^11–13^. The stability of the plasma membrane around extracellular bacteria, and its role in immune evasion and subsequent cell infection, are unknown. Ot differentiates into a distinct form prior to bacterial exit, and the intracellular (IB) and extracellular (EB) forms of Ot exhibit different protein profiles, morphology, and peptidoglycan abundance^14^. The mechanisms by which gene expression is regulated across the different stages of the infection cycle are unknown, although global transcript levels are low in the extracellular state likely reflecting its metabolically inactive state resulting from a lack of access to nutrients in the host cell cytoplasm^14^. However, the relative sense transcript levels in IB and EB form Ot are different across different genes, reflecting differentially regulated transcription between these states^14^. Relative levels of antisense transcripts across different stages of the Ot infection cycle have not been reported to date.

Ot has an unusual genome that is characterised by rampant proliferation of mobile genetic elements^15–17^. Dominant amongst these is the Rickettsial Amplified Genetic Element (RAGE) that is present in 70-90, mostly degraded, copies across the bacterial genome^18^. Whilst this RAGE is present in some species of the sister genus *Rickettsia*^19–24^, the scale of amplification is unmatched in other described organisms. It is now known that at least some Ot species encode intact, full-length RAGEs^18^ suggesting that active movement of this element may still be ongoing at a population level, but the dynamics of the Ot genome remains to be determined.

Functional antisense transcription in prokaryotes was originally described in bacteriophages and transposases, where they were shown to control the expression levels of these potentially damaging elements^25,26^. For example, *cis-*binding antisense RNA leads to degradation of the mRNA encoding a transposase gene, thus controlling transposon movement in *S. flexneri* R100 plasmid^27,28^, whilst binding of an antisense RNA at the ribosomal binding site of the anti-repressor protein Ant blocks the switch to lytic growth of *Salmonella* P22 phage^29^. More recently, pervasive antisense transcription has been described in numerous bacterial species including *E. coli, Helicobacter pylori, Bacillus subtilis, Vibrio cholerae, Chlmaydia trachomatis, Pseudomonas syringae* and *Mycobacterium tuberculosis*^26,30^. Antisense transcription can regulate genes though *cis* or *trans* activity, in which it binds to the opposite strand mRNA transcribed from the same gene, or a gene located elsewhere on the genome, respectively. *trans* active antisense small RNAs tend to interact with multiple targets, with a lower degree of sequence specificity, whilst *cis* acting antisense transcripts have perfect sequence complementarity to the opposite strand mRNA. Antisense transcripts can regulate gene expression by blocking translation or causing RNAse III dependent degradation of double stranded RNA. RNA sequencing has revealed that many bacterial genomes express antisense transcription, but it can be difficult to determine whether these result from spurious, non-specific antisense transcription initiation, or the activity of specific, cryptic antisense promoters. It has been shown that the level of genome-wide antisense transcription correlates positively with the genomic AT content^31^, which can be explained by the fact that the Pribnow motif in bacterial promoters has the consensus sequence “5’-TANAAT-3’”, so spurious promoters are much more likely in AT rich genomes. Ultimately, the biological activity of an antisense transcript can be determined experimentally through gene modification, but this is more difficult to achieve in the case of *cis* acting transcripts where the antisense transcript cannot easily be destroyed without affected the coupled sense transcript.

We previously carried out a dual RNA sequencing analysis of Ot strains Karp and UT176 grown in cultured human umbilical endothelial cells (HUVEC) in which both bacterial and host transcript abundances were determined^32^. High levels of bacterial antisense transcription were observed, with up to 1,000-fold enrichment of *cis* antisense transcripts compared to protein-coding, sense transcripts for some genes. Genes with low sense:antisense ratios were enriched in the RAGE mobile genetic element regions, and these genes were also associated with a lack of detectable peptides in a shotgun proteomics analysis carried out in parallel. One limitation of that study was that we did not independently measure antisense transcripts using alternative techniques and could not completely rule out the high levels of transcription being due to an artefact of the RNA sequencing approach used. A second limitation was that at that time the RAGE genes were not well annotated, and we therefore did not carry out a detailed analysis of the identity of RAGE genes with high antisense transcription. A new study reporting reannotation of eight Ot genomes^18^ has enabled us to perform such analysis in the current work.

Here, we sought to explore the biological significance of previously described antisense *cis* RNA transcription in Ot. We reasoned that a conserved pattern of antisense expression across different Ot strains would indicate the presence of a specific antisense promoter and potential role in bacterial growth and pathogenicity. We analysed genome-wide antisense transcription across eight related strains of Ot and found that antisense transcription is highly regulated between strains, with some genes exhibiting conserved patterns of antisense transcription along the length of the gene indicative a specific transcription start site. We carried out PCR analysis of both sense and antisense transcription of six genes in one Ot strain, the clinical isolate Karp, and found that the trends in sense:antisense ratios broadly replicated that measured by RNA sequencing. We then compared the sense:antisense ratio at different stage of the infection cycle and observed a reduction in antisense transcripts in extracellular Ot.

## Results

### High antisense transcription is conserved across diverse Ot strains

Antisense transcription has previously only been described in two Ot strains: Karp and UT176. We asked whether the presence of genes with high antisense transcription was universal across diverse Ot strains, and whether a comparative analysis could identify individual genes with consistently high antisense transcription. We carried out a dual RNA sequencing experiment in which seven Ot strains, Gilliam, Ikeda, Karp, Kato, TA686 and UT76, were grown for five days in human umbilical vein endothelial cells (HUVECs). The results of that analysis have been reported elsewhere^33^. In the current study we reanalysed this dataset, together with a previous analysis of Karp and UT176^32^ to give a total of eight different strains, with a particular focus on the distribution of genes with high levels of antisense transcription. First, we compared the levels of antisense transcription in genes encoded in the multi-copy RAGE integrative and conjugative element with those in the inter-RAGE (IR) regions (Fig. 1A). We found that all strains exhibited higher levels of antisense transcription in RAGE genes compared with IR genes, consistent with previous reports^32^. Between 1-4.3% of all inter-RAGE transcripts and 4-30% of all RAGE transcripts mapped to genes in the antisense orientation. Whilst this was a significant level of antisense transcription, it was lower than the levels reported by us previously^32^. This is likely to reflect differences in library preparation protocols, with the previous methodology specifically optimized for capturing both shorter transcripts and long (mRNA) transcripts. The relative abundance of antisense transcripts in different strains did not correlate with phylogeny^34^ or relative virulence^10^. We determined a cut-off of 200 reads to classify genes as expressing high levels of antisense transcripts and quantified the high antisense genes present in single strains or groups of strains (Fig. 1B). 18 genes were identified as having high antisense transcript levels in all 8 strains and these are listed in Fig. 1C. Of these, two encoded genes in the phage-like gene transfer agent recently identified in Ot: Karp_02209 encoding the major capsid protein and Karp_00287 encoding the tail protein. This is consistent with the described regulation of phage DNA by antisense transcription^27–29^. A comparison of the sequencing reads of these two genes across three biological replicates and eight Ot strains reveals that both genes encode a single, conserved region of high antisense transcription at the 3’ end of the gene (Fig. 1D). This high conservation suggests the presence of a specific antisense promoter in this region.

**Figure 1.**
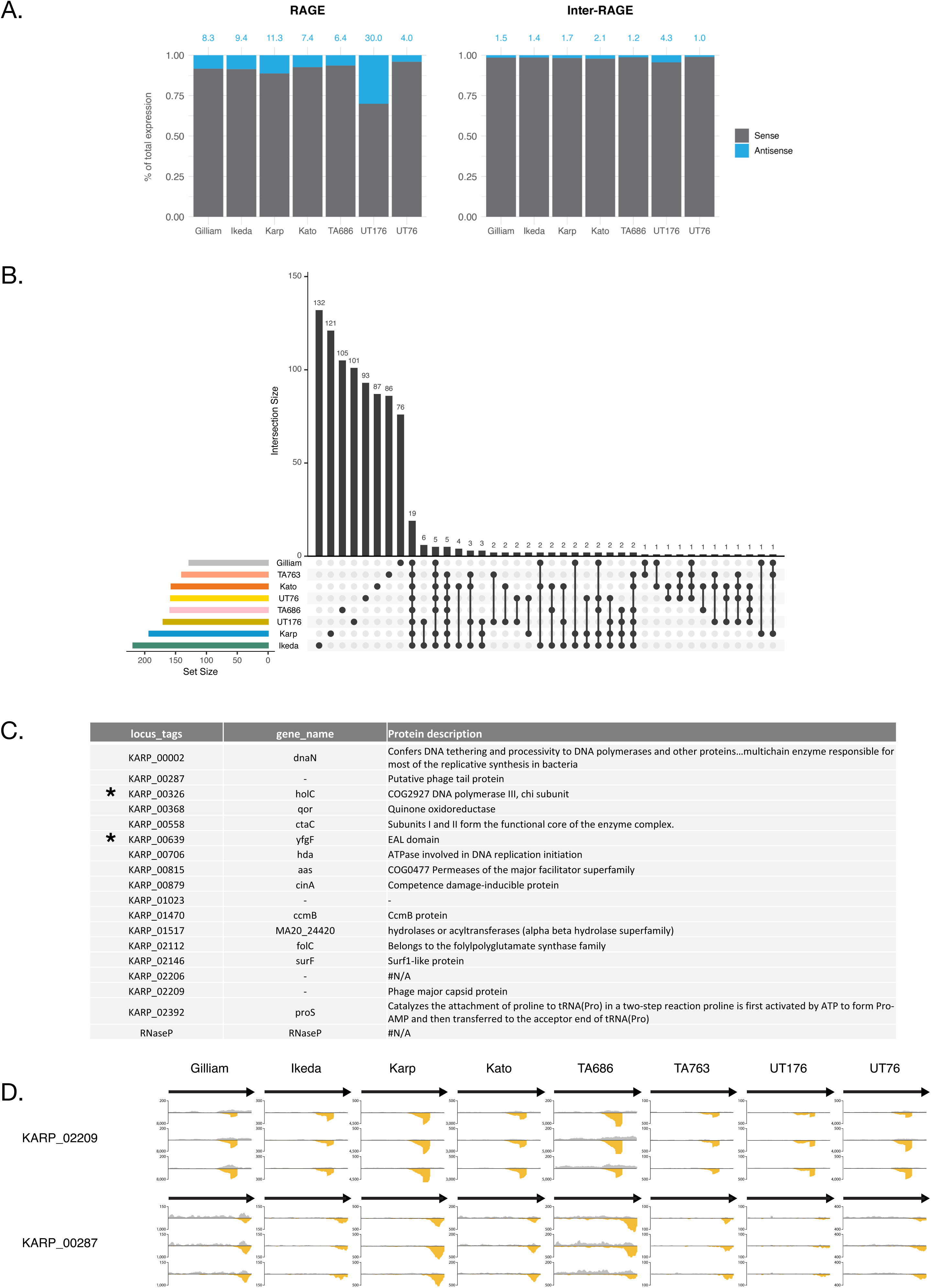
A comparison of antisense transcription across eight diverse Ot strains. An RNA sequencing analysis was carried out on eight diverse Ot strains grown in cultured human umbilical vein endothelial cells (HUVECs). A. Quantification of the % of sense and antisense transcripts from the total number of reads in each strain, separated by genes encoded in RAGE and IR regions of the bacterial genome. All strains exhibit higher antisense transcription from RAGE genes. B. A comparison of the number of high-antisense genes across genomes. Bars demonstrate the number of high-antisense genes present in individual strains and combinations of strains, as indicated by the black dots below each column. 18 genes have high antisense transcription in all eight strains. C. Description of the 18 genes with high levels of antisense transcription in all Ot strains. Asterix (*) indicates gene located within a RAGE element. D. Transcript maps showing the distribution of sense and antisense reads in the raw RNA sequencing data across three biological replicates for two gene transfer agent genes with high antisense transcripts, Karp_02209 and Karp_00287. Sense transcripts are shown in grey and antisense transcripts are shown in yellow.

### Design of a PCR assay to measure antisense transcription in Ot

We sought to validate the RNA sequencing data using an independent PCR-based experimental approach. First, we used the RNA sequencing data to select six genes with high (*ftsZ, ctrA* and *spoT)* and low (*ank12_01, folC* and *scaD)* sense:antisense transcript ratios (Fig. 2). *folC* exhibits a single antisense peak at the 3’ end of the sense transcript that is present in all strains and replicates. By contrast *ank12_01* exhibits a peak towards the middle of the gene that is not present in all strains, as well as other antisense peaks present in only some strains and replicates. *scaD* exhibits two peaks, one in the middle of the gene and one at the 3’ end of the sense strand, but these are not both universally present in all strains. Based on the RNA sequencing data we expect *folC* to exhibit the highest and most robust antisense:sense ratio.

**Figure 2.**
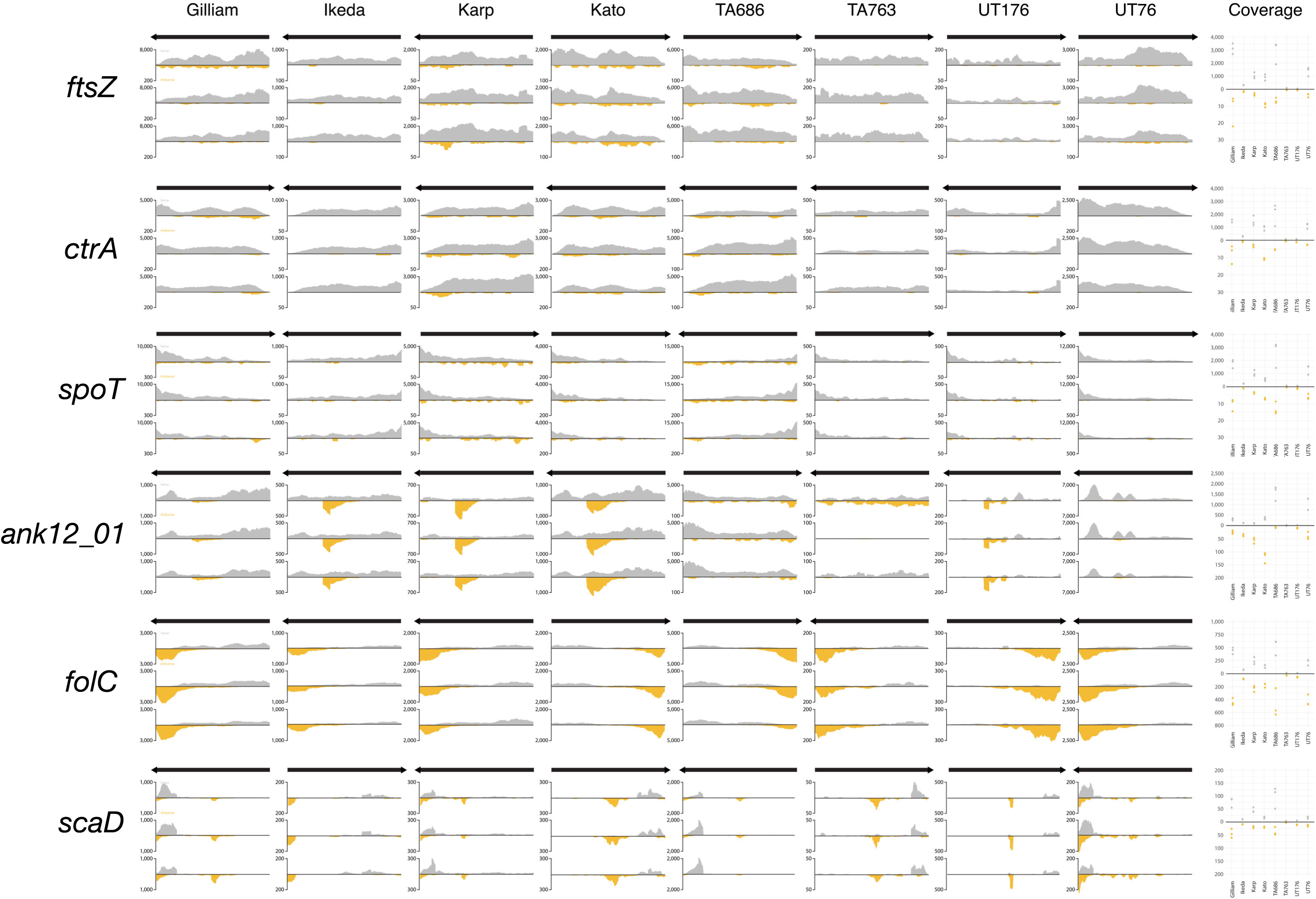
Antisense transcription across six Ot genes. Transcript maps showing the distribution of sense and antisense reads in the raw RNA sequencing data across three biological replicates for six selected Ot genes. Sense transcripts are shown in grey and antisense transcripts are shown in yellow. *ftsA, ctrA* and *spoT* exhibit low antisense transcription whilst *ank12_01, folC* and *scaD* exhibit high antisense transcription.

We designed a PCR-based assay to selectively amplify sense or antisense transcripts (Fig. 3A). We used individual primers that would only amplify sense or antisense transcripts during the reverse transcription step, followed by nested PCR amplification of cDNA. A single sense primer was designed with a binding region at the 3’ end of the gene. Since antisense transcripts do not necessarily cover the full length of the gene, we designed multiple antisense primers that would reverse transcribe RNA from different regions of the gene. The primers used for each gene are shown in Fig. 3B. In the case of genes with substantial antisense transcription predicted (Fig. 2) we designed primers covering both the specific regions with local antisense peaks and other regions across the gene.

**Figure 3.**
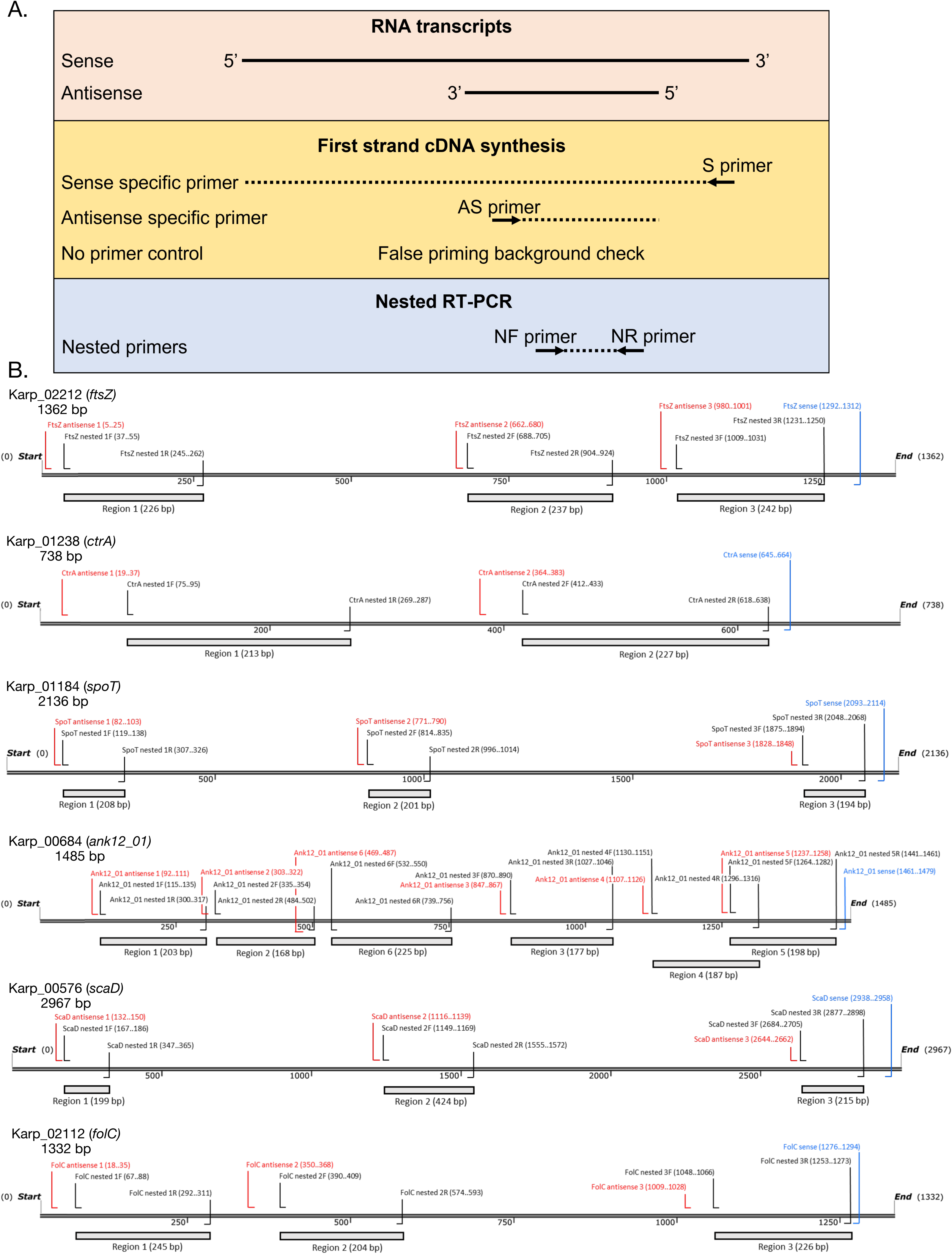
PCR assay to measure antisense transcription. A. Design of PCR assay to quantify relative levels of sense and antisense transcription. B. Gene maps showing the primers used to measure sense and antisense transcripts at multiple positions across six Ot genes.

### PCR-based quantification of sense and antisense transcripts confirms different antisense:sense ratios between six Ot genes

We used the sense:antisense PCR assay to quantify transcript levels of Ot grown in mouse fibroblast L929 cells. This was different from the human umbilical vein endothelial (HUVEC) cells used in the RNA sequencing data in order to determine the universality of our results. RNA was isolated from intracellular bacteria grown for 4 days post infection, and the relative levels of sense and antisense transcripts measured by PCR (Fig. 4A, B). We observed substantial differences between the primers designed to amplify different regions of the gene, in terms of their amplification of non-specific RNA in the no priming control as well as the relative levels of sense and antisense transcription. Broadly our results supported the predictions based on RNA sequencing data. All primer pairs against *ftsZ, ctrA* and *spoT* exhibited a sense:antisense ratio >1. For *ank12_01* and *scaD,* the genes shown to have weak and/or variable antisense peaks, the sense:antisense ratio was close to 1. Finally, for the gene *folC* that was shown to have a strong and conserved antisense peak the level of antisense was higher than the sense in all three regions tested. Region 3 corresponds to the region covering the antisense peak observed in the RNA sequencing data. Whilst this region exhibited a lower sense:antisense ratio than the other two, all three regions exhibit substantial antisense transcription.

**Figure 4.**
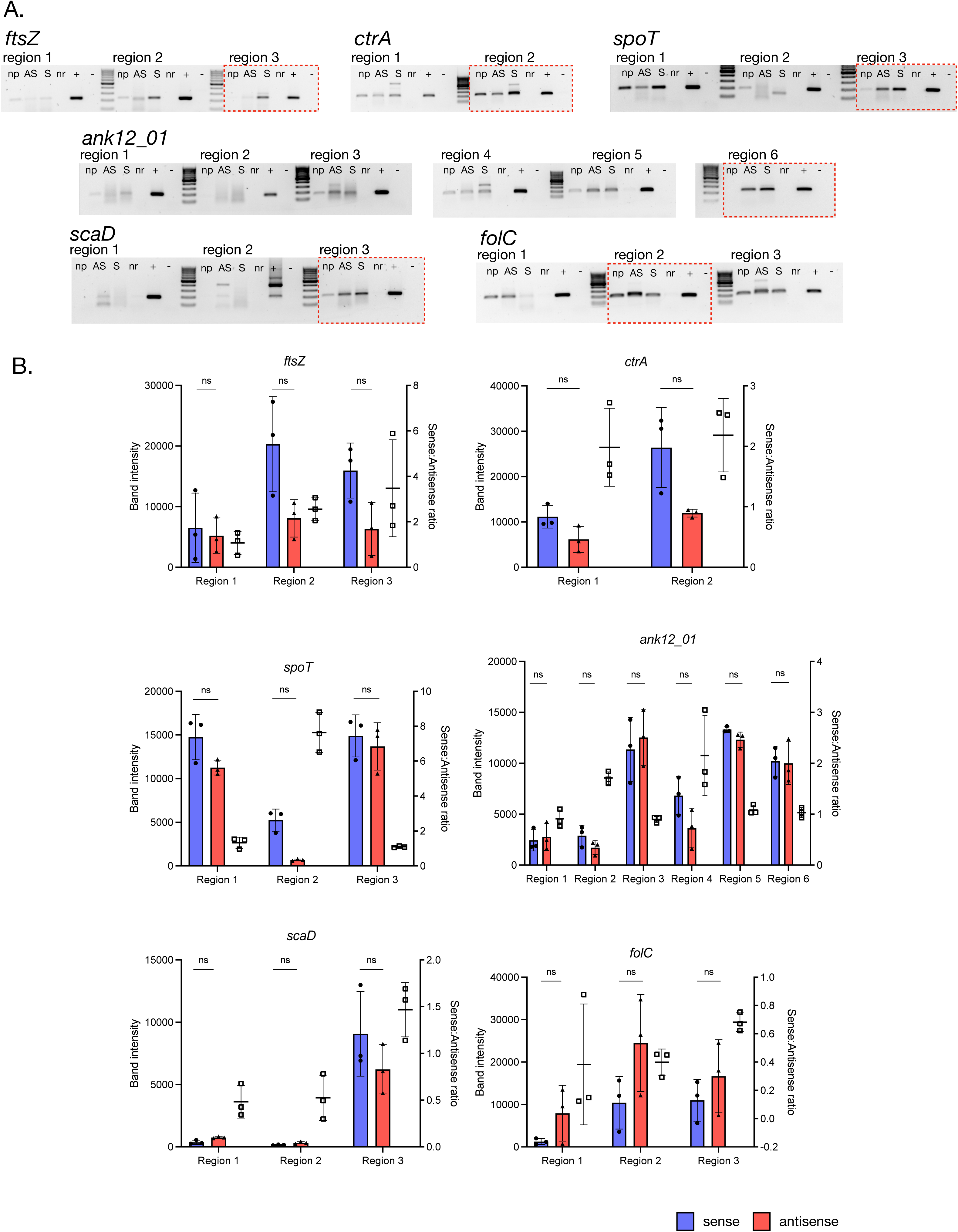
Quantification of the sense/antisense ratio at different regions within each gene. A. DNA gels showing results of PCR reactions carried out using primers shown in Fig. 3B. np = no primer used during cDNA synthesis, AS = results using antisense primer, S = results using sense primer, nr = no reverse transcriptase, + = positive control using Ot genomic DNA, - = negative control using nuclease-free water in place of DNA. 100 bp molecular weight marker was used. The region used in subsequent experiments presented in Fig. 5 are indicated with a red dotted line. B. Graphs showing quantification of data obtained in Fig. 3A. The results of three independent replicates are shown. Graph shows mean and standard deviation. Statistical significance was determined using a Mann-Whitney test where ns indicates no significant difference and * indicates p-value <0.05.

We selected one region for each gene, highlighted by a red dotted line in Fig. 4A, for subsequent analysis. These regions were selected based on the presence of antisense transcripts (where present) in RNA sequencing data and PCR analysis as well as clear amplification of a single PCR product.

### Antisense transcription is reduced in extracellular bacteria

Ot exhibits a developmental cycle in which IB form Ot is replicative and translationally active, whilst EB Ot does not replicate and is translationally inactive. Both forms are infectious. We asked whether the sense:antisense ratio would vary between these two bacterial forms. RNA was isolated from both IB and EB Ot grown in L929 cells for 5 and 7 days, and the relative levels of sense and antisense transcripts was quantified for each gene (Fig. 5). When comparing the sense and antisense reads in IB at day 5 and day 7 we observed a relatively higher level of sense transcripts compared to antisense in *ftsZ* and *ctrA,* and conversely a higher level of antisense in *folC* although this was only statistically significant on day 5. Similar ratios were observed in EB taken at 5 dpi and 7 dpi although in this case there was also a significantly higher level of sense transcripts in *spoT,* and none of the differences measured in *folC* were statistically significant. We next compared the ratios between IB and EB populations. For the high antisense genes *ank12_01, scaD* and *folA,* the sense:antisense ratios were similar between IB and EB at both time points, with the exception of day 7 where there was a significant increase in the ratio in EB compared with IB in *scaD*. By contrast, for the low antisense genes *ftsZ, ctrA* and *spoT* the levels of antisense transcription was consistently reduced in EB, resulting in an elevated sense:antisense ratio compared with IB Ot. This indicates a reduction in antisense transcription in extracellular bacteria.

**Figure 5.**
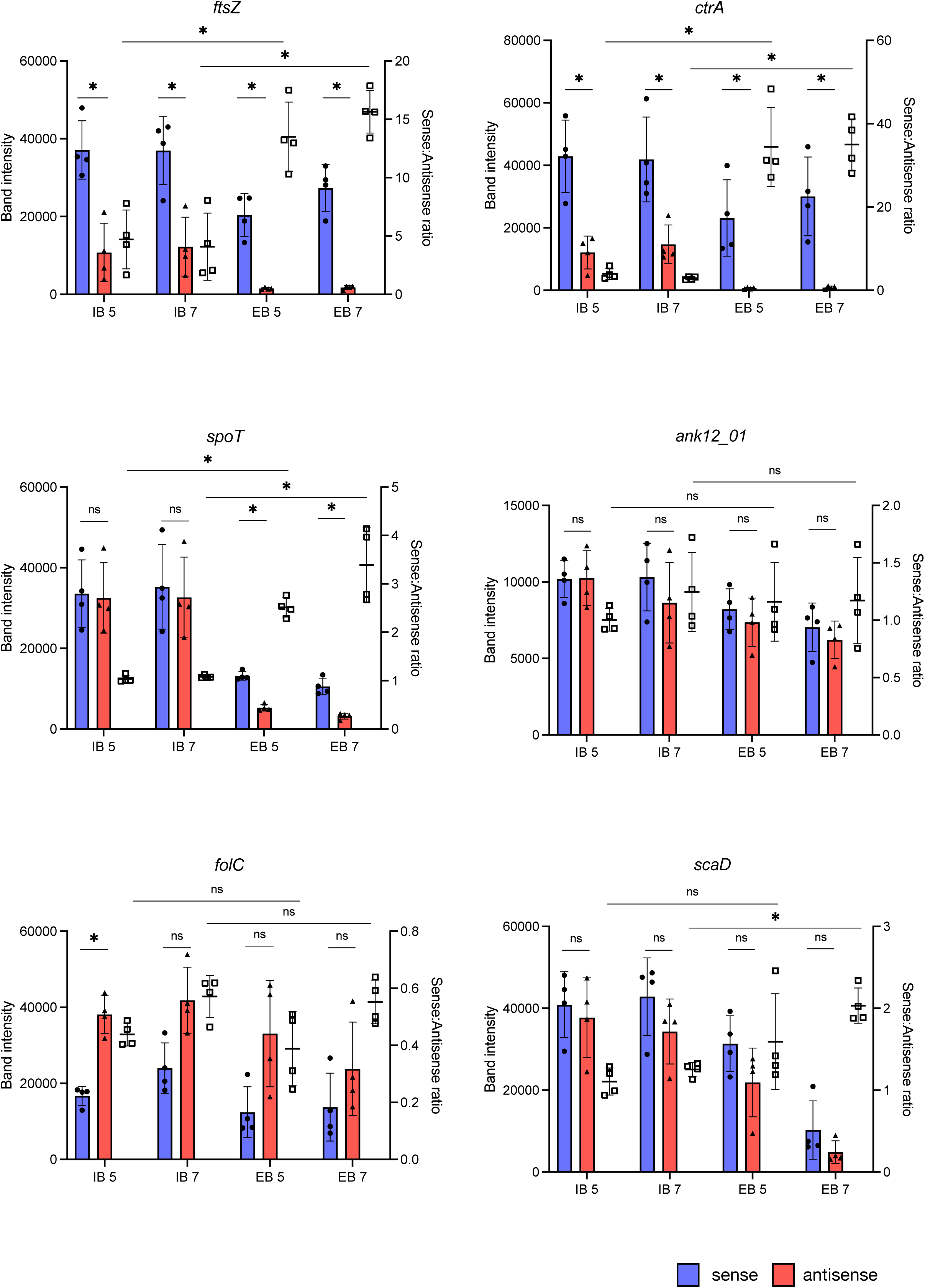
Quantification of the sense/antisense ratio at different times in the bacterial infection cycle. RNA was isolated from bacteria during rapid growth (5 days post infection) and at last stages of infection (7 days post infection). Intracellular bacteria (IB) and those that had already exited from infected cells (EB) were isolated separately. The relative levels of sense and antisense transcripts were measured for each gene at each time point. Graph shows quantification of the PCR band intensity of sense and antisense transcripts as well as the ratio of these values. The results of 3 or 4 independent biological replicates are shown, with mean and standard deviation. Statistical significance was determined using a Mann-Whitney test where ns indicates no significant difference and * indicates p-value <0.05.

## Discussion

Here we report the first detailed characterization of antisense transcription in Ot. We sought to examine whether the low sense:antisense ratio previously reported in RNA sequencing studies was a result of pervasive nonspecific transcription resulting from an AT-rich genome, or whether it was driven by specific processes in individual genes. To address this question, we carried out a large analysis of RNA sequencing data in multiple Ot strains. Examination of transcript distribution revealed genes with low sense:antisense ratios containing highly conserved antisense peaks in specific regions of the gene. This suggests the presence of a specific promoter driving antisense transcription that may inhibit gene expression through *cis-*inhibition of the coding RNA strand. It is notable that two of the genes with consistently high antisense expression across all strains encoded genes in the phage-like gene transfer agent of Ot. Whilst the activity of this agent has not been experimentally shown, its conserved presence in a highly reduced genome suggests it is functional, and it may be regulated by antisense transcription. Notably, high antisense transcription was recently reported in a related bacterium *Wolbachia,* especially in phage genes, reminiscent of our current observations^35^.

PCR quantification of the sense and antisense transcripts in different regions within a gene revealed significant differences in the magnitude of sense transcripts as well as the sense:antisense ratio. Differences in the sense transcript may result from the presence of incomplete or partially degraded transcripts, but it could also reflect differences in the efficiency of nested PCR primers for that region. Since the same nested primers are used for sense and antisense transcripts, the ratio will not be affected. *cis-*antisense transcription can initiate from multiple sites within a gene, and this is shown by the presence of specific antisense peaks in RNA sequencing reads as well as differences in the sense: antisense ratio at different regions within a gene in the PCR amplification experiment. *folC* exhibited a single, conserved antisense peak in RNA sequencing data, but this did not correlate with a single region of particularly high sense: antisense on the PCR assay. This discrepancy may be accounted for by the different cell types used, or the greater sensitivity of PCR to detect antisense transcripts compared with RNA sequencing.

The EB population of Ot is significantly different from IB. Due to its extracellular location it lacks access to essential nutrients and thus cannot replicate. Total RNA levels are extremely low and it is not known whether there remains residual transcriptional activity. The small amount of RNA present may derive from stable transcripts generated during the previous IB stage, or from ongoing, low level transcriptional activity. A comparison of the sense:antisense ratio in IB and EB revealed a further reduction in antisense transcripts in the already low antisense genes *ftsA, ctrA* and *spoT* in EB form Ot. By contrast, the sense:antisense ratio was relatively constant across IB and EB for *Ank12_01, scaD* and *folC*. The observation that antisense transcript level was reduced particularly in those genes lacking specific antisense transcription peaks in the RNA sequencing data supports the notion that the observed antisense transcripts in *Ank12_01, ScaD* and *folC* result from an active process. The general reduction of antisense transcripts in *ftsZ, ctrA* and *spoT* may reflect differential half-life of sense and antisense transcripts, with sense transcripts being specifically protected from degradation during EB stage of development.

In summary, we have shown that numerous Ot genes exhibit high antisense transcript levels, and that these are conserved between strains. This study lays the foundation for further studies on the regulation of gene expression of Ot as well as the physiology of the EB Ot population.

## Materials and Methods

### Mammalian cell culture

All experiments were performed using the mouse fibroblast cell line L929 (ECACC 85011425). The cells were grown in DMEM (Gibco 11965092) supplemented with 10% FBS (Sigma-Aldrich F7524) at 37°C and 5% CO_2_. The cells were seeded at 1.2×10^6^ cells per T75 flask two days prior to infection with bacteria.

### Bacterial propagation

L929 cells were infected with Ot strain Karp at an MOI of 200:1. The Karp infected L929 cells were then maintained at 35°C and 5% CO_2_ until the required day of harvest. Purified Karp stocks were stored in SPG solution (218 mM sucrose, 3.76 mM KH_2_PO_4_, 7.1 mM K_2_HPO_4_, and 4.9 mM potassium glutamate) at -80°C prior to use.

### Analysis of antisense transcription from RNA sequencing dataset

Orientia-infected human umbilical vein endothelial cells (HUVEC) dual RNA-seq reads were downloaded from the sequence read archive (SRA) from accession number PRJNA1047511.

### RNA-seq read processing and alignment

Quality control of sequencing reads was performed using FastQC (https://www.bioinformatics.babraham.ac.uk/projects/fastqc/) and BBDuk (https://jgi.doe.gov/data-and-tools/software-tools/bbtools/) removing low quality reads, primers and adapters.

To generate a bam file for each Ot strain, reads were aligned to a concatenated human (Gencode v38) and bacterial genome with Bowtie2^36^ using *--local* alignment, and Samtools^37^ for sorting and indexing.

### Genome coverage tracks

The bamCoverage script from DeepTools^38^ was used to generate coverage files for forward and reverse stranded reads, using parameters of *--binSize 5 --filterRNAstrand forward/reverse --outFileFormat bigwig*. The Integrative Genomics Viewer (IGV)^39^ was used to manually generate genome tracks for each gene from each of the 8 bacterial genomes, which were then combined with Adobe Illustrator. All bacterial genome and annotation files that were used in this study were taken directly from Hayward et al^33^.

### Antisense peak calling

CSaw^40^ as used to identify antisense peaks using *window size 50*, *spacing 50*, *read length 75*, *minq 10*, removing duplicated reads and excluding forward facing reads. For each strain, peaks were normalized using CPM and then identified in each of the three replicates separately (to ensure the peaks we are using are not background signals and/or artifacts). Adjacent bins from CSaw output were merged using the *reduce* function from GenomicRanges^41^. Consensus peaks were then determined if the peak was identified across all three replicates, using findOverlaps and intersect from GenomicRanges^41^. Peaks containing at least 200 reads were retained for further analysis.

### Coverage plots for Figure 2A

A custom script was created that used Samtools^37^ to extract the number of reads (coverage) for each of the specified locus tags for each strain, determining the coverage of forward and reverse reads from the three replicates.

### PCR assay

The RNA of purified Ot was extracted using Qiagen RNeasy Plus kit (Qiagen 74136) according to the manufacturer’s instructions. To avoid DNA contamination, the total RNA (5 µg) was incubated with TURBO DNase (ThermoFisher AM2238) at 37°C for 4 h before use in other PCR assays. The RNA samples were then reverse transcribed to cDNA using ImProm-II^TM^ reverse transcriptase (Promega A3802) and specific primers (sense primers and antisense primers for each target region (Supp. Table 2). cDNA synthesis was also performed on RNA samples without the addition of specific primer to detect the background amplification and on RNA samples without reverse transcriptase to ensure that there was no DNA contamination in the RNA samples. The resulting cDNA was then used as template for PCR using GoTaq^®^ DNA Polymerase (Promega M3001) and nested primers for *ftsZ*, *ctrA*, *spot*, *ank12_01*, *folC*, and *scaD*. The nested PCR products for sense and antisense expression were compared by gel electrophoresis and Gel Doc EZ System (Bio-Rad 1708270 and 1708273). The band intensities were then quantified by ImageJ software (ImageJ v1.54d).

### Preparation of IB and EB bacteria

Ot were isolated from L929 cells on 4, 5, and 7 dpi. For EB isolation, the supernatant portion of the culture was transferred to a 50 ml centrifuge tube and centrifuged at 1,000 rpm, 4°C for 5 min to remove host cells. The supernatant was then transferred into a new 50 ml centrifuge tube and centrifuged at 20,000 xg, 4°C for 15 min to collect EB pellet. For IB isolation, 1.5 ml of fresh DMEM media was added to the culture flasks. Ot infected host cells on the adhering surface were then scraped and transferred into 2 ml microcentrifuge tubes. The microcentrifuge tubes were then placed into Next Advance Bullet Blander^®^ (BBY24M-0710977) and homogenised at speed 8 for 2 min to break host cells and release IB from host cells. The cell suspensions were then centrifuged at 300 xg, RT for 3 min to remove host cells. The supernatants were then transferred into new 1.5 ml microcentrifuge tubes and centrifuged at 20,000 xg, RT for 3 min to collect IB pellet. Both IB and EB pellets were then resuspend in RNAprotect Bacteria Reagent (QIAGEN 76506) in a ratio of 2:1 (volume of RNA protect to volume of Ot pellet) and stored at -80°C until used. Hotshot was performed to quantify IB and EB DNA concentration. A portion of IB and EB cell suspension (50 ul) was pelleted and heated in 25 ul lysis buffer solution (25 mM NaOH and 0.2 mM EDTA) at 95°C for 30 min. The lysed solution is then added with 25 ul neutralization buffer (40 mM Tris-HCl) and stored at -20°C or directly used as a template in qPCR. Amplification was performed on Bio-Rad CFX96 (CT008648) using qPCRBIO Probe Mix Lo-ROX (PCR Biosystems PB20.21), 47kDa probe ([6-FAM]-TTCCACATTGTGCTGCAGATCCTTC-[TAMRA]), 47kDa forward primer (TCCAGAATTAAATGAGAATTTAGGAC), and 47kDa reverse primer (TTAGTAATTACATCTCCAGGAGCAA). The quantification of Ot copy number was then determined in relative to the 47kDa standard curve.

### Statistical analysis

The statistical analyses for band intensity were carried out using GraphPad Prism version 10.4.2 (633). A Mann-Whitney test was used to compare band intensity between two sample groups, where ns indicates no significant and * indicates p-value <0.05.

